# Microbiota Accelerates Age-Related CpG Island Methylation in Colonic Mucosa

**DOI:** 10.1101/2020.08.28.268367

**Authors:** Ang Sun, Lauren Cole, Shinji Maegawa, Pyoung Hwa Park, Kelsey Keith, Jozef Madzo, Jaroslav Jelinek, Christian Jobin, Jean-Pierre J. Issa

## Abstract

DNA methylation is an epigenetic mark that is altered in cancer and aging tissues. The effects of extrinsic factors on DNA methylation remain incompletely understood. Microbial dysbiosis is a hallmark of colorectal cancer, and infections have been linked to aberrant DNA methylation in cancers of the GI tract. To determine the microbiota’s impact on DNA methylation, we studied the methylomes of colorectal mucosa in germ-free (no microbiota) and specific-pathogen-free (controlled microbiota) mice, as well as in Il-10 KO mice (*Il10*^−/−^) which are prone to inflammation and tumorigenesis in the presence of microbiota. The presence of microbiota was associated with changes in 5% of the methylome and *Il10*^−/−^ mice showed alterations in 4.1% of the methylome. These changes were slightly more often hypo than hypermethylation and affected preferentially CpG sites located in gene bodies and intergenic regions. Mice with both Il-10 KO and microbiota showed much more pronounced alterations, affecting 18% of the methylome. When looking specifically at CpG island methylation alterations, a hallmark of aging and cancer, 0.4% were changed by the microbiota, 0.4% were changed by *Il10*^−/−^, while 4% were changed by both simultaneously. These effects are comparable to what is typically seen when comparing colon cancer to normal. We next compared these methylation changes to those seen in aging, and after exposure to the colon carcinogen Azoxymethane (AOM). Aging was associated with alterations in 18% of the methylome, and aging changes were accelerated in the *Il10*^−/−^ /SPF mice. By contrast, AOM induced profound hypomethylation that was distinct from the effects of aging or of the microbiota. CpG sites modified by the microbiota were over-represented among DNA methylation changes in colorectal cancer. Thus, the microbiota affects the DNA methylome of colorectal mucosa in patterns reminiscent of what is observed in aging and in colorectal cancer.

## Introduction

DNA methylation is an epigenetic mark with a profound impact on gene regulation and expression. This mark consists of the addition of a methyl group to a cytosine residue of a CG dinucleotide(Deaton and Bird, 2011; Issa, 2014). Approximately 80% of CpG sites in the human genome are methylated(Jelinek et al., 2012). Some CpG sites are located in CpG islands (CGIs); these islands are 500-2000 bp stretches of DNA heavily enriched for C or G bases. They remain generally unmethylated and are primarily located around transcription start sites (TSS). Methylation of TSS CGIs leads to gene silencing, but intergenic or gene body methylation can have varying effects, depending on the gene and the exact site that is methylated(Yang et al., 2014). Disruption of DNA methylation patterns is associated with aging and disease; this is characterized by global hypomethylation and aberrant hypermethylation of CGIs(Issa, 2014). CGI hypermethylation leads to the down-regulation of key genes, including tumor suppressor genes, which can directly result in tumorigenesis(Baylin and Jones, 2011; Yu et al., 2014). Consequently, studying the causes of aberrant methylation is essential to our understanding of both aging and cancer. Cell intrinsic factors such as the density of repetitive elements, baseline gene expression and binding by PCG proteins can affect the propensity to aberrant DNA methylation in cancer(Baylin and Jones, 2011; Estécio et al., 2012; Estécio et al., 2010; Estécio and Issa, 2011; Zhang et al., 2012). Cell extrinsic factors that modulate DNA methylation are less well defined, but they include aging, diet, and chronic inflammation(Chiba et al., 2012; Issa, 2014; Issa et al., 2001; Sapienza and Issa, 2016; Wallace et al., 2010).

The human gut microbiome is composed of approximately 10^13^ bacteria, which practically match the number of human cells in the body(Sender et al., 2016). The gut microbiome has co-evolved with the host, forming a symbiotic relationship contributing to energy and nutrient extraction from diets, shaping immune response, maintaining intestinal mucosal barrier integrity, and performing key xenobiotic metabolism(Holmes et al., 2012; Hooper et al., 2012; Koppel et al., 2017; Sonnenburg and Bäckhed, 2016; Turnbaugh and Gordon, 2009). The gut microbiome is linked to inflammatory diseases, a major risk factor for cancer. In addition, the microbiome has been implicated in both aging and inflammation. Germ-free mice raised and maintained without microbiota have extended lifespans relative to their normal counterparts(Tazume et al., 1991). The microbiota has the ability to induce inflammation(Tsilimigras et al., 2017) and, in turn, inflammation has been shown to significantly alter microbiota composition(Arthur et al., 2012).

Because of the common link to inflammation, there has been an interest in studying potential microbiota/epigenetic interactions. Microbiota has been shown to induce large-scale, diet-dependent changes in histone modifications(Krautkramer et al., 2016). A study of 8 pregnant women examined the microbial composition and DNA methylation in the gut, finding that CGI methylation profiles differed based on microbial composition(Kumar et al., 2014). Additionally, infections have been linked to DNA methylation changes in cancer: Infection with *H. pylori* led to methylation of CGI promoters including TSGs in gastric cancer(Maekita et al., 2006). Also in gastric cancer, a hypermethylation phenotype termed CpG Island Methylator Phenotype (CIMP) has been linked to Epstein-Barr virus (EBV) presence(Chang et al., 2006). It was also shown that a specific family of bacteria, *Fusobacteria,* is markedly enriched in colorectal cancers (CRCs) with CIMP, but much less so in cancers without CIMP(Tahara et al., 2014b). Based on these data, we hypothesized that the microbiota could induce changes in DNA methylation. To test this hypothesis directly, we compared DNA methylomes in the intestines of germ-free (GF) mice and mice inoculated with microbiota that was specific-pathogen-free (SPF). We also studied interactions between the presence of microbiota and the methylomes of Il-10 KO mice (prone to inflammation and tumorigenesis), aging mice, and mice exposed to the carcinogen Azoxymethane (AOM).

## Methods

Mouse tissues for microbiota, *Il10*^−/−^ and AOM studies were obtained from previously described experiments testing the effects of microbiota on colonic tumors*(Arthur et al., 2012)*. Germ-free (GF) *Il10*^−/−^ and WT mice (129SvEv) were raised in sterile conditions and kept free of microbiota. Specific-pathogen-free (SPF) mice were kept in controlled conditions and free of pathogenic microbes. Mouse tissues for aging studies were obtained from 6 mice kept in sterile conditions on regular diets and sacrificed at 4 and 30-33 months of age. We studied a total of 48 mice overall (Table 1).

**Table 1:** Type and number of mice studied for DNA methylation

| Mouse Model | Number of Mice | Comment |
| --- | --- | --- |
| WT/GF | 6 | Germ-free (GF) mice – no microbiota |
| WT/SPF | 6 | Mice with a Specific-Pathogen-Free (SPF) microbiota |
| *Il10*<sup>-/-</sup>/GF | 6 | *Il10*<sup>-/-</sup> are predisposed to spontaneous inflammation but only in the presence of microbiota |
| *Il10*<sup>-/-</sup>/SPF | 6 |  |
| WT/SPF + AOM | 6 | Azoxymethane (AOM) is a colon carcinogen |
| *Il10*<sup>-/-</sup>/GF + AOM | 6 | Predisposed to spontaneous inflammation and tumorigenesis but only in the presence of microbiota |
| *Il10*<sup>-/-</sup>/SPF + AOM | 6 |  |
| C57/B6 WT | 6 | Ages 4, 4, 4, 30, 33 and 33 months |

We performed Digital Restriction Enzyme Analysis of Methylation (DREAM) on DNA extracted from mouse proximal colon tissues, as described^3^. DREAM is a quantitative, deep-sequencing based method of measuring DNA methylation at CpG sites within the CCCGGG sequence. DNA is sequentially treated with two restriction enzymes: *Sma*I and *Xma*I, which act on the same DNA sequence, but leave different 5’ ends of fragments. *SmaI* is blocked by CpG methylation, but *XmaI* is not. By treating DNA first with *SmaI,* then *XmaI*, we created distinct signatures for unmethylated and methylated CpG sites at the edges of restriction fragments^3^. Next, we ligated Illumina sequencing adapters to the ends of the restriction fragments and sequenced the resulting libraries on Illumina HiSeq 2500 instrument at Fox Chase Cancer Center Genomics facility. We aligned the sequencing reads to the mouse genome (mm9) using Bowtie2, and counted signatures corresponding to unmethylated and methylated CpGs. We calculated methylation levels as the ratio of reads with the methylated signature to all reads mapping to each respective site. We adjusted DNA methylation values at individual CpG sites based on spiked in standards and filtered for the minimum sequencing depth of 100 reads.

The datasets generated during this study are available at GEO (accession number: GSE150333, https://www.ncbi.nlm.nih.gov/geo/query/acc.cgi?acc=GSE150333).

Statistical analyses were performed, and figures were generated using R(R Core Team, 2019) (retrieved from https://www.R-project.org/). Volcano plots were generated by plotting the average difference in methylation for a given CpG site against the negative log10 of the p-value. Average methylation change was calculated by subtracting average methylation in one condition from another, and an unpaired t-test was used to calculate a p-value. Bar plots were generated by taking the number of sites meeting a condition dividing by the number of sites examined. For each CpG site, we generated a multivariate linear regression for the relationship between methylation and microbiota, Il-10 deficiency, and AOM treatment. The linear equation is defined by y = b_m_x_m_ + b_i_x_i_ + b_a_x_a_ + b_0_ + ε, where y is methylation, x is the respective condition (m = microbiota, i = inflammation, or a = azoxymethane respectively), and b is the respective regression coefficient. UpSet plots were generated using the ggupset package(Ahlmann-Eltze, 2020), while all other plots were generated using the ggplot2 package. All reported p-values are two-sided, and p≤0.05 was used as a threshold of significance.

## Results

### The microbiota modulates DNA methylation

We used DREAM to examine how the presence of microbiota affects DNA methylation. To do this, DREAM data for wild-type (WT) germ-free (GF) and wild-type specific-pathogen-free (SPF) mice were analyzed and a volcano plot depicting the methylation differences between them was generated (Figure 1a). For most experiments, each group of mice consisted of 6 animals (Table 1). DREAM detected the methylation status of 24,865 sites on average at a minimum of 10 reads/site. We considered sites to be “changed” if there was a statistically significant increase or decrease in average methylation of 5% or more. To ensure precision, we only analyzed CpG sites that had greater than 100 reads in at least 75% of the mice in each group. Overall, of 12,919 detectable sites, 3.7% decreased, and 1.3% of sites increased in methylation. Thus, 5% of CpG sites analyzed showed differences triggered by the presence of microbiota.

**Figure 1:**
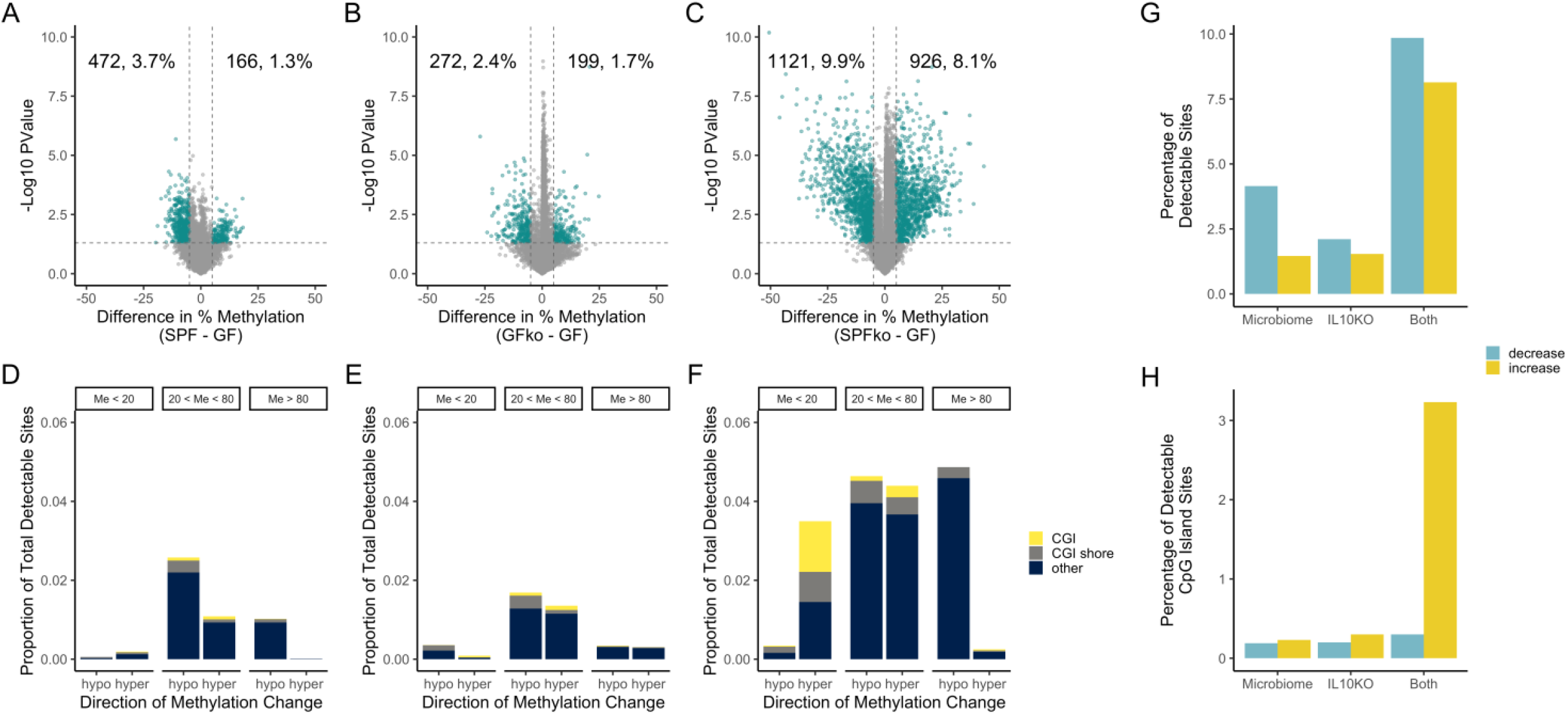
microbiota influences DNA methylation. A-C) Volcano plot analysis showing methylation differences between SPF and GF mice (A), *Il10*^−/−^ and GF mice (B), and SPF-*Il10*^−/−^ and GF mice (C). The x-axis shows the difference in average methylation between SPF and GF mice for a given site. The y-axis is the negative log(10) of the p-value, which was determined with a Student’s t-test. All sites above the dotted line are significant at p≤0.05. Green sites change at a magnitude of 5% or greater. D-F) Bar graphs showing the proportion and type of CpG sites that change at least 5% between SPF and GF mice (D), *Il10*^−/−^ and GF mice (E), and SPF-*Il10*^−/−^ and GF mice (F). Shore indicates sites that are not in CGIs, but within 2000 bp of them. (G) Bar graph showing the proportion of CpG sites that show DNA methylation alterations in the volcano plots in A-C. (H) Bar graph showing the proportion of CpG sites within CpG islands that show DNA methylation alterations in the volcano plots in A-C.

Sites that are greater than 80% or less than 20% methylated at baseline tend to be more stable than intermediately methylated sites(Jelinek et al., 2012). Therefore, we examined how the microbiota affect different CpG compartments. Volcano plots were generated that looked only at sites with greater than 80% average methylation, sites with less than 20% methylation and sites with between 20 and 80 percent methylation in GF mice (Figure S1). Overall, 0.3% of unmethylated sites were affected by microbiota, compared to 1.1% of highly methylated sites, and 3.8% of intermediately methylated sites. Figure 1d shows the distribution of sites that were changed at least 5% between GF and SPF mice, stratified by baseline methylation. CGI and CGI shore sites appear to be largely stable. The most vulnerable sites seem to be gene body or intergenic areas with baseline methylation of between 20% and 80%. Thus, the presence of microbiota significantly affects DNA methylation in colonic mucosa with slightly more pronounced hypomethylation than hypermethylation.

### Deletion of *Il10* affects DNA methylation

Interleukin 10 (*Il10*) is an anti-inflammatory cytokine and mice lacking this gene develop spontaneous intestinal inflammation in the presence of microbiota(Sellon et al., 1998). To determine the effects of Il10 on intestinal DNA methylation independent of microbiota, DREAM data from GF/WT and GF/*Il10*^−/−^ mice were analyzed; Figure 1b shows the volcano plot. Of all detectable sites, 2.4% decreased and 1.7% increased in methylation. Figure 1e shows the distribution of affected sites located in CGIs, CGI shores or other sites; Like Figure 1d, Figure 1e shows that relatively few CGI/shore sites were subject to change, and most of the changes that did occur were at variable sites with 20% to 80% methylation (also see Figure S1). GF/*Il10*^−/−^ mice have little detectable inflammation; thus, the DNA methylation changes seen in these animals may be direct effects of Il10 deficiency or may be related to low grade/patchy inflammation that is not detectable using the usual assays.

Il10 deficiency combined with microbiota results in inflammation and markedly accelerates colon tumorigenesis in mice(Uronis et al., 2009). By comparing DREAM data from GF/WT to SPF/*Il10*^−/−^ mice, we were able to observe the effects of both conditions occurring simultaneously. In a differential methylation analysis (Figure 1e), we found that 8.1% of sites increased and 9.9% of sites decreased in average methylation. In addition to more sites changing average methylation, the combined effects of inflammation susceptibility and microbiota appear to cause a specific increase in methylation at CGIs and CGI shores. Figure 1f shows that 2% of all sites were CGI/CGI shore sites that increased in average methylation between WT/GF and SPF/*Il10*^−/−^ mice. This contrasted with the individual effects of IL10 deficiency or microbiota, which had little impact on CGIs or CGI shores. Figure 1g gives a broad overview of the effects of microbiota and *Il10*^−/−^ on DNA methylation individually and together. Both microbiota and inflammation changed methylation at approximately 4-5% of sites. When combined, approximately 18% of sites experienced average methylation changes, with more sites decreasing in average methylation than increasing. When we examined CGIs exclusively (Figure 1h), 0.4% were changed by SPF, 0.5% were changed by *Il10*^−/−^ while 3.5% were changed by both simultaneously. These dramatic differences in DNA methylation are comparable to what can be seen when comparing cancer to normal(Maegawa et al., 2010).

### Differential effects of azoxymethane and *Il10*^−/−^ on DNA methylation

Azoxymethane (AOM) is an agent commonly used to induce tumorigenesis in mouse models of colorectal cancer. We sought to determine if AOM and *Il10*^−/−^ differed in their effects on DNA methylation profiles. We examined the effects of AOM alone, *Il10*^−/−^ alone, and both in combination in the presence of microbiota (i.e. in SPF mice). A differential methylation analysis showed that AOM had a pronounced hypomethylating effect (Figure 2a). Approximately 8.1% of sites decreased in average methylation as a result of AOM, as opposed to only 1.2% of sites increasing in average methylation. It is interesting to note that in both sites with 80% or more average methylation and 20% or less average methylation, AOM caused overall decreases in average methylation (Figure 2b). In addition, AOM did not seem to target CGI or CGI shore sites and was most likely to cause a decrease in methylation of sites with medium levels of baseline methylation (20%<Me<80%, Figure 2b). By contrast, *Il10*^−/−^ in SPF mice led to 5.7% of sites increasing in methylation and 6.2% of sites decreasing in methylation (Figure 2C). About 2% of sites were located in CGIs or CGI shores that increased in methylation (Figure 2D). Thus, AOM and *IL10*^−/−^ both induced hypomethylation but, in the presence of microbiota, only *Il10*^−/−^ induced substantial CGI hypermethylation, pointing to potentially different mechanisms for their effects on DNA methylation.

**Figure 2:**
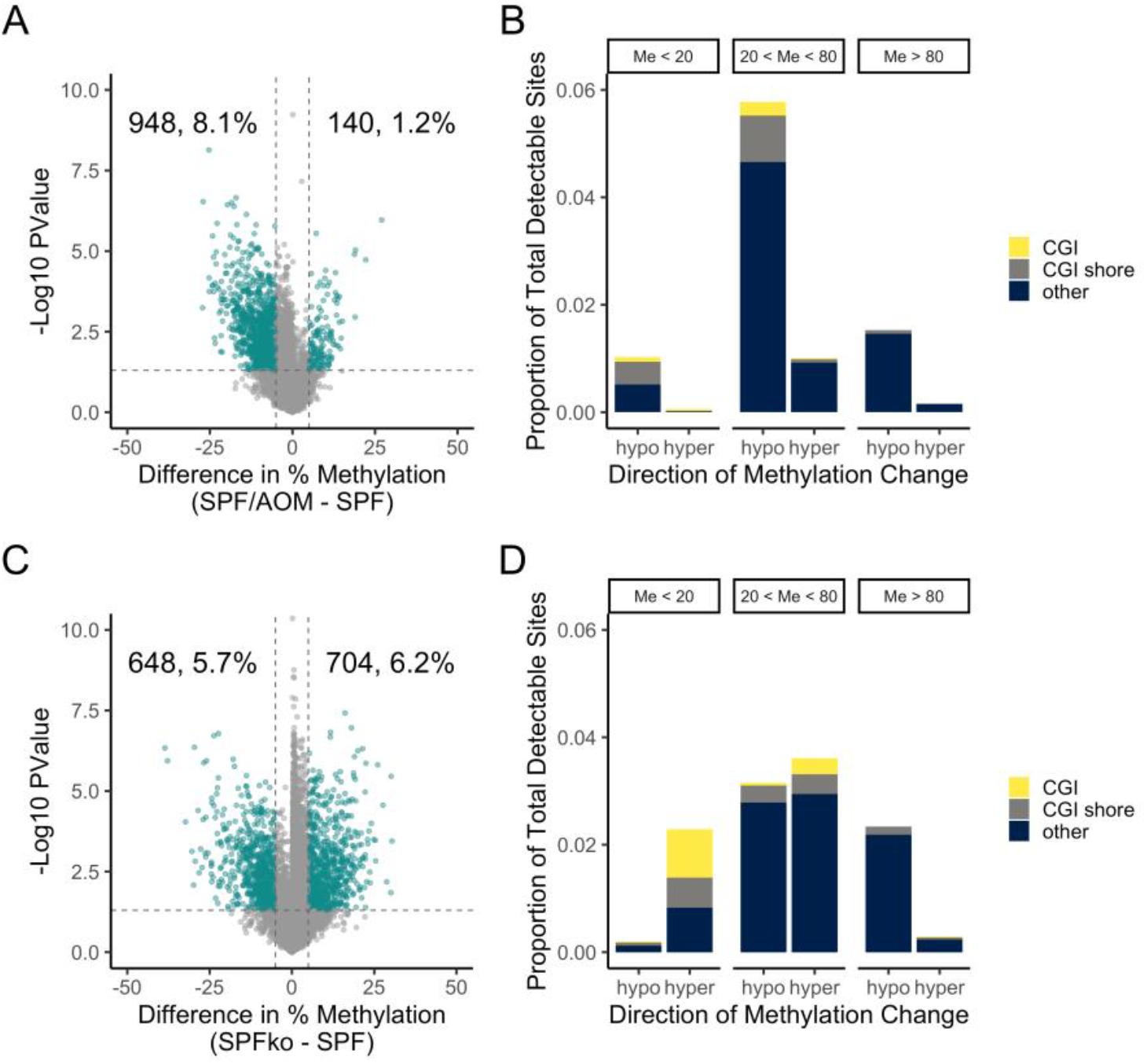
AOM is a potent hypomethylating carcinogen. (A) Volcano plot analysis showing methylation differences between SPF and SPF+AOM mice. See Figure 1 for graph details. (B) Bar graph showing the proportion and type of CpG sites that change at least 5% in (A). (C) Volcano plot analysis showing methylation differences between SPF and SPF-*Il10*^−/−^ mice. See Figure 1 for graph details. (D) Bar graph showing the proportion and type of CpG sites that change at least 5% in (C).

### Shared and unique DNA methylation changes

With multiple simultaneous treatments, it is not possible to determine in a simple differential methylation test whether the methylation is affected by a single treatment or a combination of treatments. To elucidate and compare the individual effects of an SPF microbiota, Il10 deficiency, and AOM treatment on DNA methylation, we built an additive linear regression model that incorporates data from all 42 mice studied. Using an FDR of 0.05, we found that the linear model recapitulated the trends seen in the individual comparisons. Using this model (Figure 3a), we determined that microbiota (SPF) induced changes in 13.5% of sites (6.9% hyper and 6.5% hypomethylated), and Il10 deficiency induced changes in 10.5% of sites (9% hyper and 1.5% hypomethylated), while AOM induced changes in 5.9% of sites (1% hyper and 4.9% hypomethylated). AOM continued to show a pronounced hypomethylating effect, while *Il10*^−/−^ continued to show a strong hypermethylating effect. Microbiota show equal hypomethylating and hypermethylating effects. We used UpSet plots to determine the overlap in effects of the different exposures on CpG methylation, examined separately for hypomethylation (Fig. 3b) and hypermethylation (Fig. 3c). Microbiota and *Il10*^−/−^ have the most shared events, particularly when it came to hypermethylation. Thus, of 865 sites hypermethylated by the microbiota, 671 (78%) were also affected by *Il10*^−/−^. There was less conservation when it came to hypomethylation (22% of sites affected by the microbiota were also affected by *Il10*^−/−^). AOM had mostly unique effects. These data suggest that the microbiota and *Il10*^−/−^ have shared methylome interaction mechanisms, while AOM has an independent mechanism of action.

**Figure 3:**
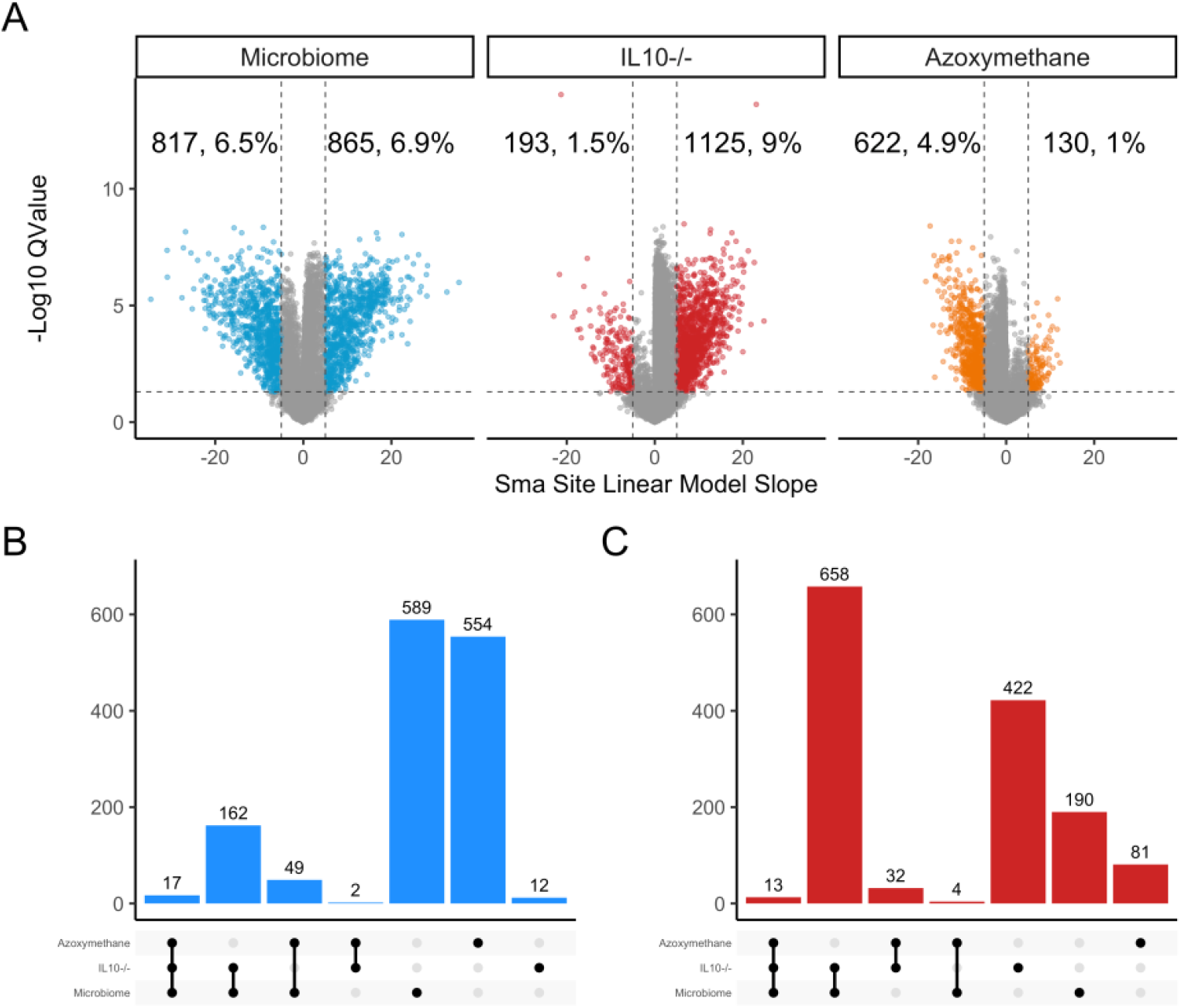
Linear regression model of the effects of microbiota, *Il10*^−/−^ and AOM on DNA methylation. (A) Shows Volcano plots of FDR (0.05) corrected significant methylation changes attributed to each external exposure. The x-axis indicates the linear model slope for individual CpG sites (equivalent to % change in methylation) while the y-axis is the negative log(10) of the q-value. (B) Shows Upset plots of shared/unique hypomethylation events while (C) shows UpSet plots of shared/unique hypermethylation events. The number of shared or uniquely altered CpG sites is indicated on top of each vertical bar.

### The microbiota and inflammation accelerate age-related methylation drift

Methylation drift characterized by a global loss of DNA methylation with simultaneous hypermethylation in CGI promoters occurs as a result of aging(Maegawa et al., 2014; Maegawa et al., 2017). To compare this to microbiota effects, we first generated DREAM data for old (aged 30 months) and young (aged 4-5 months) mouse colon (Figure 4a). Looking at aging in C57/Bl6 wild type mice, we found that of ~20,000 detectable sites, 6.6% decreased in methylation, and 11.5% increased in methylation. Hypermethylated CGIs accounted for 7% of the changes. We analyzed the overlap between sites that changed in methylation due to aging and sites that changed due to microbiota, Il10 deficiency or AOM. Aging had the largest effect individually but there were many shared hypomethylation (Figure 4b) and hypermethylation (Figure 4c) events between aging and the extrinsic exposures. For example, out of 472 sites hypomethylated upon the microbiota exposure, 139 (29%) were also affected by age; of the 272 sites affected by *Il10*^−/−^, 31% were also affected by age; of the 1121 sites hypomethylated in inflamed mice (SPF-*Il10*^−/−^), 31% were hypomethylated in aging mice. Similarly, out of 166 sites hypermethylated upon the microbiota exposure, 18% were also affected by age; of the 199 sites affected by *Il10*^−/−^, 33% were also affected by age; of the 926 sites hypomethylated in inflamed mice (SPF-*Il10*^−/−^), 35% were also hypermethylated in aging mice. This corroborates previous data demonstrating partial overlap between inflammation-related and age-related methylation(Hahn et al., 2008), and implicates the microbiota in this process.

**Figure 4:**
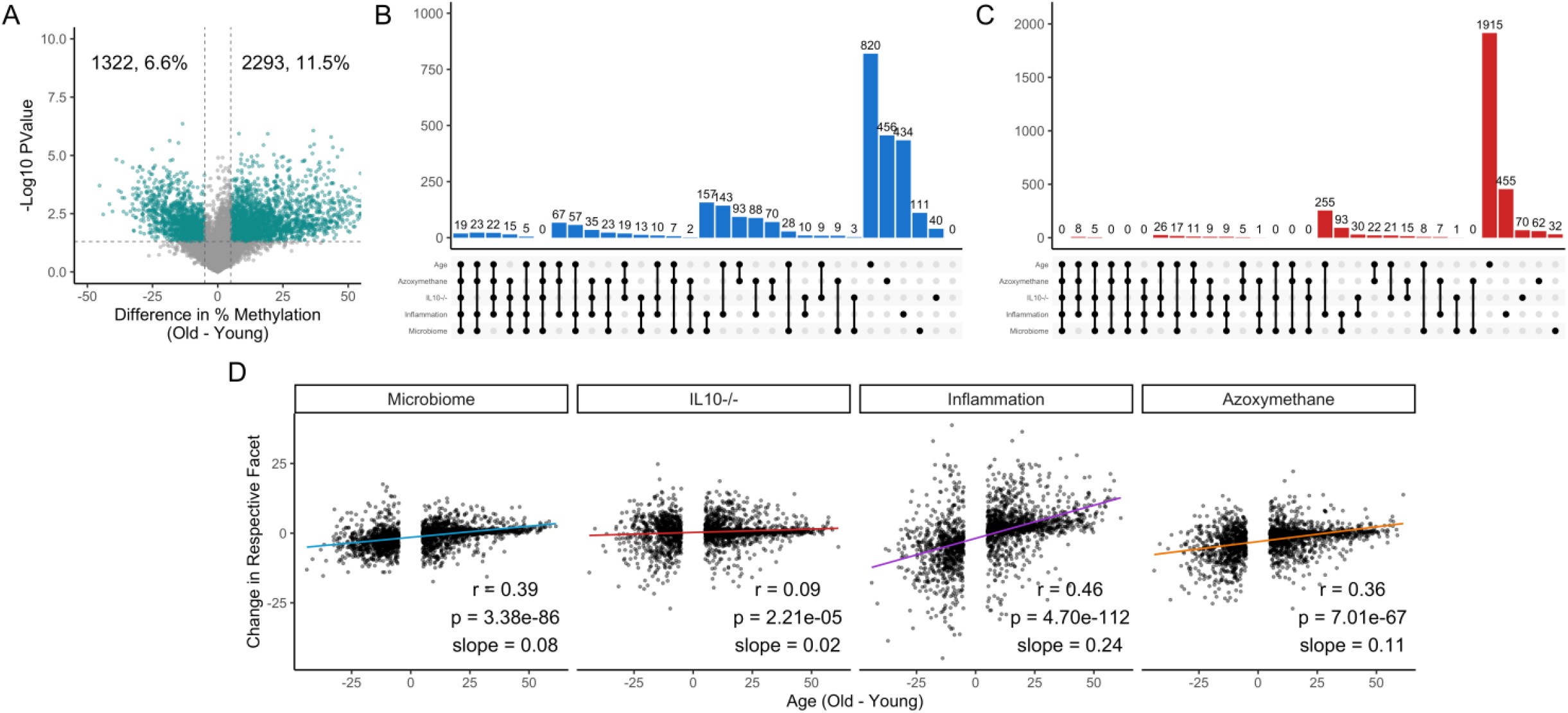
microbiota and inflammation modify the same CpG sites subject to age-related methylation drift. (A) Volcano plot analysis showing methylation differences between young and old mice. See Figure 1A for graph details. (B) UpSet plots of shared/unique hypomethylation events between sites that decrease at least 5% during aging and in the different exposures analyzed in Figures 1-3. “Inflammation” refers to *Il10*^−/−^/SPF mice. (C) Same analysis as in B for sites that increase methylation at least 5%. (D) Scatterplot of average methylation change with age (x-axis) to average change by exposures (y-axis) for all sites that change at least 5% with age. Pearson r, p-value, and slope are indicated in each plot.

One drawback to this analysis is the use of a threshold of 5% change in average methylation to consider a site affected by inflammation, microbiota, or aging; this threshold could lead to underestimation of conservation because of sites that may change at a lower magnitude in one condition. To address this, we focused on the large number of sites that change with age and generated scatterplots comparing a site’s average change with age to change caused by exposures. Interestingly, the correlation coefficients were positive and statistically significant for all exposures with the strongest correlations for inflammation (SPF-*Il10*^−/−^, r=0.46, p<0.0001) followed by microbiota (r=0.39, p<0.0001), AOM exposure (r=.36, p<0.0001) and IL10 deficiency (r=0.09, p<0.0001). Taken together, these data indicate that the vast majority of sites that change with aging were affected by microbiota/inflammation combination. There were also sites that were affected by exposures but not by aging (Figure S4), most evident in mice with inflammation triggered by the combined effects of SPF and *IL10*^−/−^.

### CpG sites affected by the microbiota are hypermethylated in colon cancer

To determine the impact of microbiota, inflammation, and/or aging on methylation in colon cancer, we analyzed data from The Cancer Genome Atlas (TCGA). To correspond mouse to human data, we focused on promoters (defined as −1500 to 500 of transcription start site), and we used data on the CpG site closest to the transcription start site. Converging DREAM data with TCGA data, we were able to analyze 2,042 genes in total. Figure 5a shows scatter plots of methylation changes in colon cancer patients included in the TCGA data set (β value of methylation difference between tumor and normal) plotted on the y-axis, and methylation changes in our datasets (based on the volcano plot for aging and on the linear model for individual exposures) plotted on the x-axis. The plots demonstrate a strong concordance between methylation changes in TCGA and each of aging, microbiota exposure, and IL10 deficiency. We calculated odds ratios to quantitate this concordance, at a threshold of 5% change in TCGA and in the extrinsic exposures. For hypermethylation, the odds ratios for enrichment were 15.5 (95% CI 10.6-22.9; q = 3.0×10^−54^) for age, 0 (95% CI 0-0; q = 1) for azoxymethane, 2.3 (95% CI 1.1-4.5; q = 0.039) for *IL10*^−/−^, and 4.1 (95% CI 1.8-9.0; q = 9.4×10^−4^) for microbiota. For hypomethylation, odds ratios were 2.0 (95% CI 0.04-13.9; q = 0.47) for age, 9.4 (95% CI 1.6-38.9; q = 0.015) for azoxymethane, 9.2 (95% CI 0.2-116.2; q = 0.18) for *IL10*^−/−^ and 11.5 (95% CI 2.6-41.1; q = 0.003) for microbiota. Thus, as previously reported, aging has a major impact on whether a gene becomes hypermethylated in cancer, but we also find that genes affected by microbiota and by Il10 deficiency are also overrepresented among genes hypermethylated in CRC.

**Figure 5:**
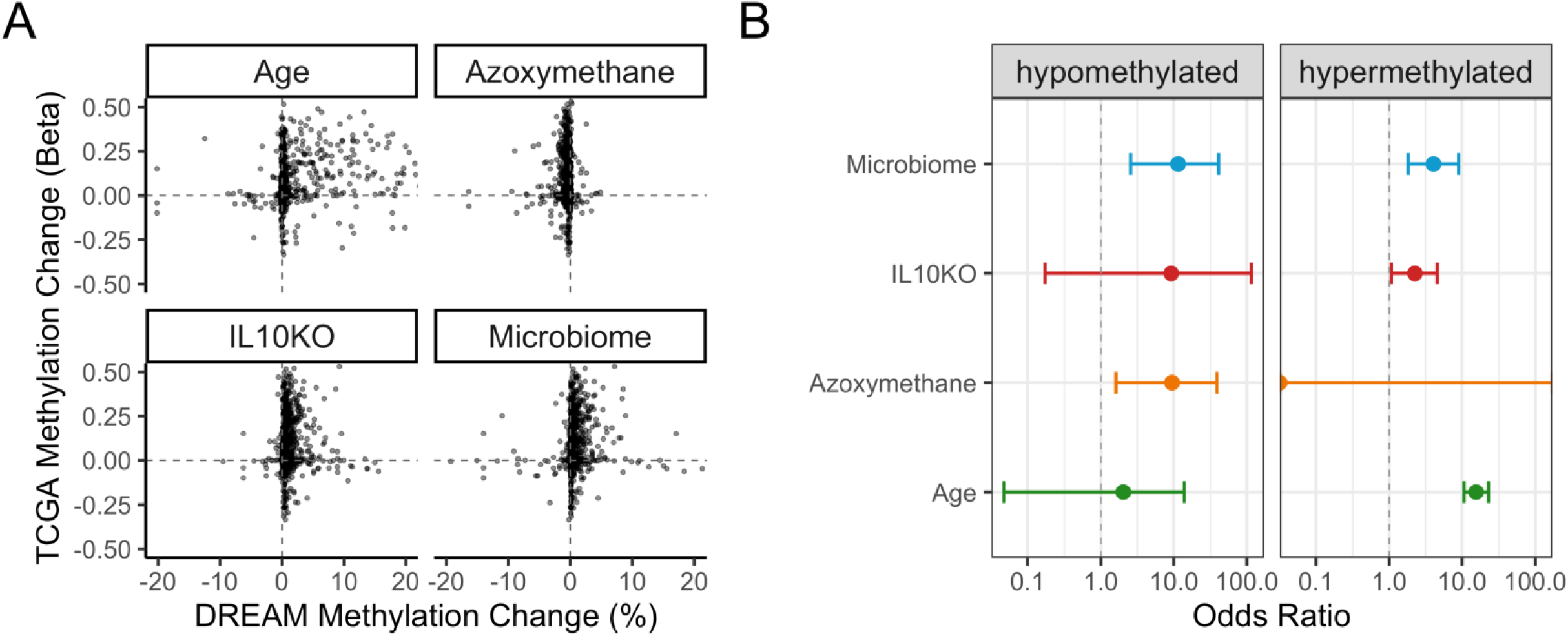
Genes affected by extrinsic exposures are more likely to be altered in cancer. (A) Scatter plot of DNA methylation change by aging or exposure (x-axis) compared to DNA methylation change in colon cancer TCGA samples. (B) Odds ratios compute enrichment of genes altered by aging or different extrinsic exposures among genes altered in colon cancer (TCGA data). The odds ratios are computed separately for gene promoters hypomethylated (left) or hypermethylated (right) by exposures and in colon cancer.

## Discussion

We show here that the presence of microbiota causes changes in DNA methylation that affect 5% of the methylome, a high number when one considers the fact that DNA methylation is very carefully controlled with relatively low levels of inter-individual variability(Jelinek et al., 2012). These effects are compounded by the presence of inflammation – mice with both Il-10 deficiency and exposure to the microbiota showed alterations in 18% of the methylome, and the differences were particularly striking at CpG islands, regions that are normally very stable and highly protected from DNA methylation(Bird, 2002). This strikingly high number is quantitatively similar to what can be seen when comparing mice at the extremes of their lifespan(Maegawa et al., 2010), and the pattern of change is very similar to what can be seen in colon cancers (CGI hypermethylation, intergenic hypomethylation). Indeed, CpG sites affected by the microbiota are overrepresented among genes hypermethylated in colon cancer, suggesting that microbiota could have an important influence on shaping cancer epigenomes, as previously suggested(Tahara et al., 2014b). Interestingly, while the presence of microbiota accelerates age-related methylation drift, there are also epigenetic changes specific to the different extrinsic factors studied (microbiota, inflammation, AOM) suggesting distinct mechanisms rather than non-specific effects of cycles of injury and stem cell proliferation. It is also worth noting that the effects we observe are related to pathogen-free microbiota. It would be interesting to determine whether more profound changes are seen when pathogenic bacteria are introduced into the mix.

Our results are consistent with recent studies but extend the findings substantially. Three studies reported that microbiota affects the physiology and DNA methylation patterns in developing intestinal epithelium(Ansari et al., 2020; Pan et al., 2018; Yu et al., 2015). Interestingly, the effects seen were primarily outside promoters (e.g. 3’ end of genes and intergenic areas) while our results highlight the profound effect of microbiota and inflammation on promoter CpG islands – the compartment most uniquely affected in aging and cancer(Issa, 2014). Ansari also studied the effects of inflammation induced by Dextran Sodium Sulfate on DNA methylation, and found primarily hypomethylation(Ansari et al., 2020), in marked contrast to what we observed with inflammation induced by the combination of Il-10 deficiency and the presence of microbiota. Dextran Sodium Sulfate’s effects are reminiscent of what we observed with exposure to azoxymethane and raise the possibility that the effects seen are related to the chemical’s effect more directly than to inflammation. Finally, Sobhani et al. introduced into germ-free mice human fecal microbiota from patients with colon cancer or controls and also observed alterations in DNA methylation(Sobhani et al., 2019), and some of the genes affected (such as *SFRP1*) are known to show age-related methylation in human colon(Shen et al., 2007). In our study, we show that even a pathogen-free microbiota affects DNA methylation at relatively high levels. Our data uniquely highlight the effects of microbiota and inflammation on aberrant CpG island DNA methylation and link it directly to the acceleration of aging effects on DNA methylation. The data also suggest that microbiota/inflammation effects are potential precursors to the aberrant DNA methylation seen in colorectal cancers.

The mechanism by which microbiota affects DNA methylation remains to be elucidated. One possibility is via the production of metabolites which act as cofactors or inhibitors for epigenetic enzymes. For example, the microbial metabolites acetate, propionate, and butyrate are capable of inducing mass reprogramming of histone methylation and acetylation states(Krautkramer et al., 2016). Bacterial species producing these short-chain fatty acid metabolites are known to be decreased in cancer and inflammation. DNA methyltransferases (DNMTs) have been shown to interact with modifications on histone tails(Baubec and Schübeler, 2014), and as histone states change, DNA methylation patterns may as well. Additionally, the gut bacteria may cause alterations in host gene expression that affect the ability of host cells to uptake certain metabolites; for example, it has been demonstrated that *E. coli* is capable of downregulating a protein responsible for butyrate uptake(Kumar et al., 2015). Aberrant DNA methylation may also be directly initiated by the overproduction of oncogenic metabolites (oncometabolites). Loss of function mutations in citric acid cycle enzymes fumarate hydratase (FH) and succinate dehydrogenase (SDH) result in the accumulation of fumarate and succinate respectively(Yang et al., 2013). A neomorphic mutation in isocitrate dehydrogenase (IDH) causes the enzyme to produce 2-hydroxyglutarate (2HG) instead of α-ketoglutarate(Ward et al., 2010). Each of these mutations is found in certain types of CIMP positive cancer(Figueroa et al., 2010), though they are very rarely observed in colorectal cancer(Tahara et al., 2014a). 2HG binds to and inhibits α-ketoglutarate dependent TET and methyltransferase enzymes, causing DNA hypermethylation and tumorigenesis(Figueroa et al., 2010). Succinate and fumarate have been shown to inhibit these enzymes *in-vitro* and there is overlap between the methylated genes associated with each metabolite anomaly, implying all three may act via this same mechanism(Yang et al., 2013). Thus, it is plausible that bacteria cause aberrant methylation in part via over-production of metabolites that modulate the function of the TET DNA demethylases.

Previous studies also showed that inflammation alters the DNA methylome(Issa, 2014; Niwa et al., 2010), though the precise mechanisms by which this occurs is unknown. It has been proposed that inflammation increases cell turnover rate, and inflammation-induced methylation may simply be a reflection of an increased rate of age-related methylation(Sapienza and Issa, 2016). The overlap we observed between inflammation and aging-associated changes are consistent with this. However, we found that microbiota and inflammation also induced methylation changes that were not observed in aging. It has been suggested that direct interactions with inflammatory cytokines alter a cell’s methylation profile(Gasche et al., 2011). It is also possible that the inflamed gut exerts different selective pressures on host intestinal epithelial cells than the non-inflamed gut, selecting for cells with unique DNA methylation profiles. It may eventually be possible to tease out inflammation-independent effects of extrinsic factors on DNA methylation.

In conclusion, we find that microbiota has a profound effect on DNA methylation in the gut and helps shape the aberrant methylomes seen in aging and cancer. The fact that extrinsic factors (bacteria, inflammation, carcinogens) modulate the epigenome suggests potential ways to intervene therapeutically for cancer prevention.

## Acknowledgments

This work was supported by National Institutes of Health grants R01-CA214005.

## Author Contributions

Conceptualization, A.S. and J.J.I.; Methodology, A.S. and J.J.I.; Formal Analysis, A.S., L.C., S.M., P.H.P, K.K., J.M., and J.J.; Investigation, A.S.; Resources, C.J.; Data Curation, A.S., S.M., and J.J; Writing – Original Draft, A.S., L.C., and J.J.I.; Writing – Review & Editing, A.S., C.J., and J.J.I.; Visualization, A.S., L.C., and J.J.I.; Supervision, J.J.I.; Project Administration, A.S. and J.J.I; Funding Acquisition, J.J.I.

## References

Ahlmann-Eltze, C. (2020). ggupset: Combination Matrix Axis for ‘ggplot2’ to Create ‘UpSet’ Plots. R package version 0.3.0. DOI: https://CRAN.R-project.org/package=ggupset.

Ansari, I., Raddatz, G., Gutekunst, J., Ridnik, M., Cohen, D., Abu-Remaileh, M., Tuganbaev, T., Shapiro, H., Pikarsky, E., Elinav, E., et al. (2020). The microbiota programs DNA methylation to control intestinal homeostasis and inflammation. Nat Microbiol. 5(4), 610–619. Published online 2020/02/03 DOI: 10.1038/s41564-019-0659-3.

Arthur, J.C., Perez-Chanona, E., Mühlbauer, M., Tomkovich, S., Uronis, J.M., Fan, T.J., Campbell, B.J., Abujamel, T., Dogan, B., Rogers, A.B., et al. (2012). Intestinal inflammation targets cancer-inducing activity of the microbiota. Science. 338(6103), 120–123. DOI: 10.1126/science.1224820.

Baubec, T., and Schübeler, D. (2014). Genomic patterns and context specific interpretation of DNA methylation. Curr Opin Genet Dev. 25, 85–92. Published online 2014/03/07 DOI: 10.1016/j.gde.2013.11.015.

Baylin, S.B., and Jones, P.A. (2011). A decade of exploring the cancer epigenome - biological and translational implications. Nat Rev Cancer. 11(10), 726–734. DOI: 10.1038/nrc3130.

Bird, A. (2002). DNA methylation patterns and epigenetic memory. Genes Dev. 16(1), 6–21. DOI: 10.1101/gad.947102.

Chang, M.S., Uozaki, H., Chong, J.M., Ushiku, T., Sakuma, K., Ishikawa, S., Hino, R., Barua, R.R., Iwasaki, Y., Arai, K., et al. (2006). CpG island methylation status in gastric carcinoma with and without infection of Epstein-Barr virus. Clin Cancer Res. 12(10), 2995–3002. DOI: 10.1158/1078-0432.CCR-05-1601.

Chiba, T., Marusawa, H., and Ushijima, T. (2012). Inflammation-associated cancer development in digestive organs: mechanisms and roles for genetic and epigenetic modulation. Gastroenterology. 143(3), 550–563. Published online 2012/07/13 DOI: 10.1053/j.gastro.2012.07.009.

Deaton, A.M., and Bird, A. (2011). CpG islands and the regulation of transcription. Genes Dev. 25(10), 1010–1022. DOI: 10.1101/gad.2037511.

Estécio, M.R., Gallegos, J., Dekmezian, M., Lu, Y., Liang, S., and Issa, J.P. (2012). SINE retrotransposons cause epigenetic reprogramming of adjacent gene promoters. Mol Cancer Res. 10(10), 1332–1342. DOI: 10.1158/1541-7786.MCR-12-0351.

Estécio, M.R., Gallegos, J., Vallot, C., Castoro, R.J., Chung, W., Maegawa, S., Oki, Y., Kondo, Y., Jelinek, J., Shen, L., et al. (2010). Genome architecture marked by retrotransposons modulates predisposition to DNA methylation in cancer. Genome Res. 20(10), 1369–1382. DOI: 10.1101/gr.107318.110.

Estécio, M.R., and Issa, J.P. (2011). Dissecting DNA hypermethylation in cancer. FEBS Lett. 585(13), 2078–2086. DOI: 10.1016/j.febslet.2010.12.001.

Figueroa, M.E., Abdel-Wahab, O., Lu, C., Ward, P.S., Patel, J., Shih, A., Li, Y., Bhagwat, N., Vasanthakumar, A., Fernandez, H.F., et al. (2010). Leukemic IDH1 and IDH2 mutations result in a hypermethylation phenotype, disrupt TET2 function, and impair hematopoietic differentiation. Cancer Cell. 18(6), 553–567. DOI: 10.1016/j.ccr.2010.11.015.

Gasche, J.A., Hoffmann, J., Boland, C.R., and Goel, A. (2011). Interleukin-6 promotes tumorigenesis by altering DNA methylation in oral cancer cells. Int J Cancer. 129(5), 1053–1063. Published online 2011/01/07 DOI: 10.1002/ijc.25764.

Hahn, M.A., Hahn, T., Lee, D.H., Esworthy, R.S., Kim, B.W., Riggs, A.D., Chu, F.F., and Pfeifer, G.P. (2008). Methylation of polycomb target genes in intestinal cancer is mediated by inflammation. Cancer Res. 68(24), 10280–10289. DOI: 10.1158/0008-5472.CAN-08-1957.

Holmes, E., Li, J.V., Marchesi, J.R., and Nicholson, J.K. (2012). Gut microbiota composition and activity in relation to host metabolic phenotype and disease risk. Cell Metab. 16(5), 559–564. DOI: 10.1016/j.cmet.2012.10.007.

Hooper, L.V., Littman, D.R., and Macpherson, A.J. (2012). Interactions between the microbiota and the immune system. Science. 336(6086), 1268–1273. Published online 2012/06/06 DOI: 10.1126/science.1223490.

Issa, J.P. (2014). Aging and epigenetic drift: a vicious cycle. J Clin Invest. 124(1), 24–29. DOI: 10.1172/JCI69735.

Issa, J.P., Ahuja, N., Toyota, M., Bronner, M.P., and Brentnall, T.A. (2001). Accelerated age-related CpG island methylation in ulcerative colitis. Cancer Res. 61(9), 3573–3577.

Jelinek, J., Liang, S., Lu, Y., He, R., Ramagli, L.S., Shpall, E.J., Estecio, M.R., and Issa, J.P. (2012). Conserved DNA methylation patterns in healthy blood cells and extensive changes in leukemia measured by a new quantitative technique. Epigenetics. 7(12), 1368–1378. DOI: 10.4161/epi.22552.

Koppel, N., Maini Rekdal, V., and Balskus, E.P. (2017). Chemical transformation of xenobiotics by the human gut microbiota. Science. 356(6344). DOI: 10.1126/science.aag2770.

Krautkramer, K.A., Kreznar, J.H., Romano, K.A., Vivas, E.I., Barrett-Wilt, G.A., Rabaglia, M.E., Keller, M.P., Attie, A.D., Rey, F.E., and Denu, J.M. (2016). Diet-Microbiota Interactions Mediate Global Epigenetic Programming in Multiple Host Tissues. Mol Cell. 64(5), 982–992. Published online 2016/11/23 DOI: 10.1016/j.molcel.2016.10.025.

Kumar, A., Alrefai, W.A., Borthakur, A., and Dudeja, P.K. (2015). Lactobacillus acidophilus counteracts enteropathogenic E. coli-induced inhibition of butyrate uptake in intestinal epithelial cells. Am J Physiol Gastrointest Liver Physiol. 309(7), G602–607. Published online 2015/08/13 DOI: 10.1152/ajpgi.00186.2015.

Kumar, H., Lund, R., Laiho, A., Lundelin, K., Ley, R.E., Isolauri, E., and Salminen, S. (2014). Gut microbiota as an epigenetic regulator: pilot study based on whole-genome methylation analysis. MBio. 5(6). Published online 2014/12/16 DOI: 10.1128/mBio.02113-14.

Maegawa, S., Gough, S.M., Watanabe-Okochi, N., Lu, Y., Zhang, N., Castoro, R.J., Estecio, M.R., Jelinek, J., Liang, S., Kitamura, T., et al. (2014). Age-related epigenetic drift in the pathogenesis of MDS and AML. Genome Res. 24(4), 580–591. DOI: 10.1101/gr.157529.113.

Maegawa, S., Hinkal, G., Kim, H.S., Shen, L., Zhang, L., Zhang, J., Zhang, N., Liang, S., Donehower, L.A., and Issa, J.P. (2010). Widespread and tissue specific age-related DNA methylation changes in mice. Genome Res. 20(3), 332–340. DOI: 10.1101/gr.096826.109.

Maegawa, S., Lu, Y., Tahara, T., Lee, J.T., Madzo, J., Liang, S., Jelinek, J., Colman, R.J., and Issa, J.J. (2017). Caloric restriction delays age-related methylation drift. Nat Commun. 8(1), 539. Published online 2017/09/14 DOI: 10.1038/s41467-017-00607-3.

Maekita, T., Nakazawa, K., Mihara, M., Nakajima, T., Yanaoka, K., Iguchi, M., Arii, K., Kaneda, A., Tsukamoto, T., Tatematsu, M., et al. (2006). High levels of aberrant DNA methylation in Helicobacter pylori-infected gastric mucosae and its possible association with gastric cancer risk. Clin Cancer Res. 12(3 Pt 1), 989–995. DOI: 10.1158/1078-0432.CCR-05-2096.

Niwa, T., Tsukamoto, T., Toyoda, T., Mori, A., Tanaka, H., Maekita, T., Ichinose, M., Tatematsu, M., and Ushijima, T. (2010). Inflammatory processes triggered by Helicobacter pylori infection cause aberrant DNA methylation in gastric epithelial cells. Cancer Res. 70(4), 1430–1440. DOI: 10.1158/0008-5472.CAN-09-2755.

Pan, W.H., Sommer, F., Falk-Paulsen, M., Ulas, T., Best, P., Fazio, A., Kachroo, P., Luzius, A., Jentzsch, M., Rehman, A., et al. (2018). Exposure to the gut microbiota drives distinct methylome and transcriptome changes in intestinal epithelial cells during postnatal development. Genome Med. 10(1), 27. Published online 2018/04/13 DOI: 10.1186/s13073-018-0534-5.

R Core Team, 2019. R: A language and environment for statistical computing. R Foundation for Statistical Computing, https://www.R-project.org, Vienna, Austria.

Sapienza, C., and Issa, J.P. (2016). Diet, Nutrition, and Cancer Epigenetics. Annu Rev Nutr. 36, 665–681. Published online 2016/03/23 DOI: 10.1146/annurev-nutr-121415-112634.

Sellon, R.K., Tonkonogy, S., Schultz, M., Dieleman, L.A., Grenther, W., Balish, E., Rennick, D.M., and Sartor, R.B. (1998). Resident enteric bacteria are necessary for development of spontaneous colitis and immune system activation in interleukin-10-deficient mice. Infect Immun. 66(11), 5224–5231. DOI: 10.1128/IAI.66.11.5224-5231.1998.

Sender, R., Fuchs, S., and Milo, R. (2016). Revised Estimates for the Number of Human and Bacteria Cells in the Body. PLoS Biol. 14(8), e1002533. Published online 2016/08/19 DOI: 10.1371/journal.pbio.1002533.

Shen, L., Toyota, M., Kondo, Y., Lin, E., Zhang, L., Guo, Y., Hernandez, N.S., Chen, X., Ahmed, S., Konishi, K., et al. (2007). Integrated genetic and epigenetic analysis identifies three different subclasses of colon cancer. Proc Natl Acad Sci U S A. 104(47), 18654–18659. DOI: 10.1073/pnas.0704652104.

Sobhani, I., Bergsten, E., Couffin, S., Amiot, A., Nebbad, B., Barau, C., de’Angelis, N., Rabot, S., Canoui-Poitrine, F., Mestivier, D., et al. (2019). Colorectal cancer-associated microbiota contributes to oncogenic epigenetic signatures. Proc Natl Acad Sci U S A. 116(48), 24285–24295. Published online 2019/11/11 DOI: 10.1073/pnas.1912129116.

Sonnenburg, J.L., and Bäckhed, F. (2016). Diet-microbiota interactions as moderators of human metabolism. Nature. 535(7610), 56–64. DOI: 10.1038/nature18846.

Tahara, T., Yamamoto, E., Madireddi, P., Suzuki, H., Maruyama, R., Chung, W., Garriga, J., Jelinek, J., Yamano, H.O., Sugai, T., et al. (2014a). Colorectal carcinomas with CpG island methylator phenotype 1 frequently contain mutations in chromatin regulators. Gastroenterology. 146(2), 530–538.e535. DOI: 10.1053/j.gastro.2013.10.060.

Tahara, T., Yamamoto, E., Suzuki, H., Maruyama, R., Chung, W., Garriga, J., Jelinek, J., Yamano, H.O., Sugai, T., An, B., et al. (2014b). Fusobacterium in colonic flora and molecular features of colorectal carcinoma. Cancer Res. 74(5), 1311–1318. DOI: 10.1158/0008-5472.CAN-13-1865.

Tazume, S., Umehara, K., Matsuzawa, H., Aikawa, H., Hashimoto, K., and Sasaki, S. (1991). Effects of germfree status and food restriction on longevity and growth of mice. Jikken Dobutsu. 40(4), 517–522.

Tsilimigras, M.C., Fodor, A., and Jobin, C. (2017). Carcinogenesis and therapeutics: the microbiota perspective. Nat Microbiol. 2, 17008. Published online 2017/02/22 DOI: 10.1038/nmicrobiol.2017.8.

Turnbaugh, P.J., and Gordon, J.I. (2009). The core gut microbiome, energy balance and obesity. J Physiol. 587(Pt 17), 4153–4158. Published online 2009/06/02 DOI: 10.1113/jphysiol.2009.174136.

Uronis, J.M., Mühlbauer, M., Herfarth, H.H., Rubinas, T.C., Jones, G.S., and Jobin, C. (2009). Modulation of the intestinal microbiota alters colitis-associated colorectal cancer susceptibility. PLoS One. 4(6), e6026. DOI: 10.1371/journal.pone.0006026.

Wallace, K., Grau, M.V., Levine, A.J., Shen, L., Hamdan, R., Chen, X., Gui, J., Haile, R.W., Barry, E.L., Ahnen, D., et al. (2010). Association between folate levels and CpG Island hypermethylation in normal colorectal mucosa. Cancer Prev Res (Phila). 3(12), 1552–1564. DOI: 10.1158/1940-6207.CAPR-10-0047.

Ward, P.S., Patel, J., Wise, D.R., Abdel-Wahab, O., Bennett, B.D., Coller, H.A., Cross, J.R., Fantin, V.R., Hedvat, C.V., Perl, A.E., et al. (2010). The common feature of leukemia-associated IDH1 and IDH2 mutations is a neomorphic enzyme activity converting alpha-ketoglutarate to 2-hydroxyglutarate. Cancer Cell. 17(3), 225–234. Published online 2010/02/18 DOI: 10.1016/j.ccr.2010.01.020.

Yang, M., Soga, T., and Pollard, P.J. (2013). Oncometabolites: linking altered metabolism with cancer. J Clin Invest. 123(9), 3652–3658. Published online 2013/09/03 DOI: 10.1172/JCI67228.

Yang, X., Han, H., De Carvalho, D.D., Lay, F.D., Jones, P.A., and Liang, G. (2014). Gene body methylation can alter gene expression and is a therapeutic target in cancer. Cancer Cell. 26(4), 577–590. DOI: 10.1016/j.ccr.2014.07.028.

Yu, D.H., Gadkari, M., Zhou, Q., Yu, S., Gao, N., Guan, Y., Schady, D., Roshan, T.N., Chen, M.H., Laritsky, E., et al. (2015). Postnatal epigenetic regulation of intestinal stem cells requires DNA methylation and is guided by the microbiome. Genome Biol. 16, 211. Published online 2015/09/30 DOI: 10.1186/s13059-015-0763-5.

Yu, D.H., Waterland, R.A., Zhang, P., Schady, D., Chen, M.H., Guan, Y., Gadkari, M., and Shen, L. (2014). Targeted p16(Ink4a) epimutation causes tumorigenesis and reduces survival in mice. J Clin Invest. 124(9), 3708–3712. Published online 2014/07/25 DOI: 10.1172/JCI76507.

Zhang, Y., Shu, J., Si, J., Shen, L., Estecio, M.R., and Issa, J.P. (2012). Repetitive elements and enforced transcriptional repression co-operate to enhance DNA methylation spreading into a promoter CpG-island. Nucleic Acids Res. 40(15), 7257–7268. DOI: 10.1093/nar/gks429.

