## Supplementary Figures for "Microbiota Accelerates Age-Related CpG Island Methylation in Colonic Mucosa"

Figure S1: microbiota influences DNA methylation, stratified by baseline methylation. (A) Volcano plots show methylation differences between GF and SPF mice, analyzing sites with less than 20% methylation on average (left), sites with between 20% and 80% methylation (middle) and sites with greater than 80% methylation (right). (B) Similar analyses of methylation differences between *Il10*$^{-/-}$ and GF mice.

Figure S4: microbiota and inflammation modify the same CpG sites subject to age-related methylation drift. Shown are scatterplots similar to Figure 4(D) but with reversed axes, with average methylation change with age (y-axis) to average change by exposures (x-axis) for all sites that change at least 5% with exposures. Pearson R, p-value and slope are indicated in each plot.

### Figure S1

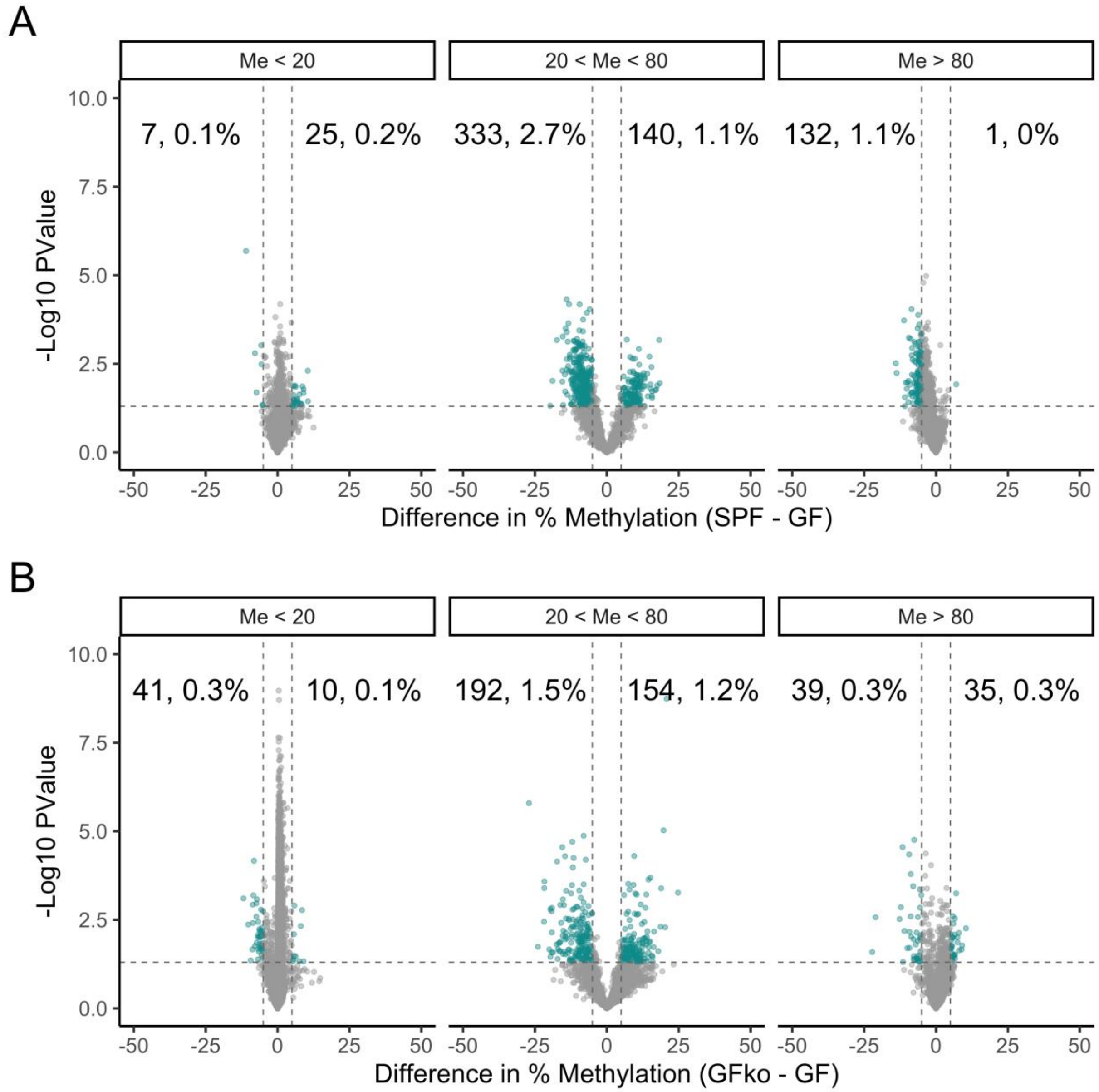
A
Me < 20
20 < Me < 80
Me > 80
7, 0.1%
25, 0.2%
333, 2.7%
140, 1.1%
132, 1.1%
1, 0%
-Log10 PValue
Difference in % Methylation (SPF - GF)
B
Me < 20
20 < Me < 80
Me > 80
41, 0.3%
10, 0.1%
192, 1.5%
154, 1.2%
39, 0.3%
35, 0.3%
-Log10 PValue
Difference in % Methylation (GFko - GF)

### Figure S4

A

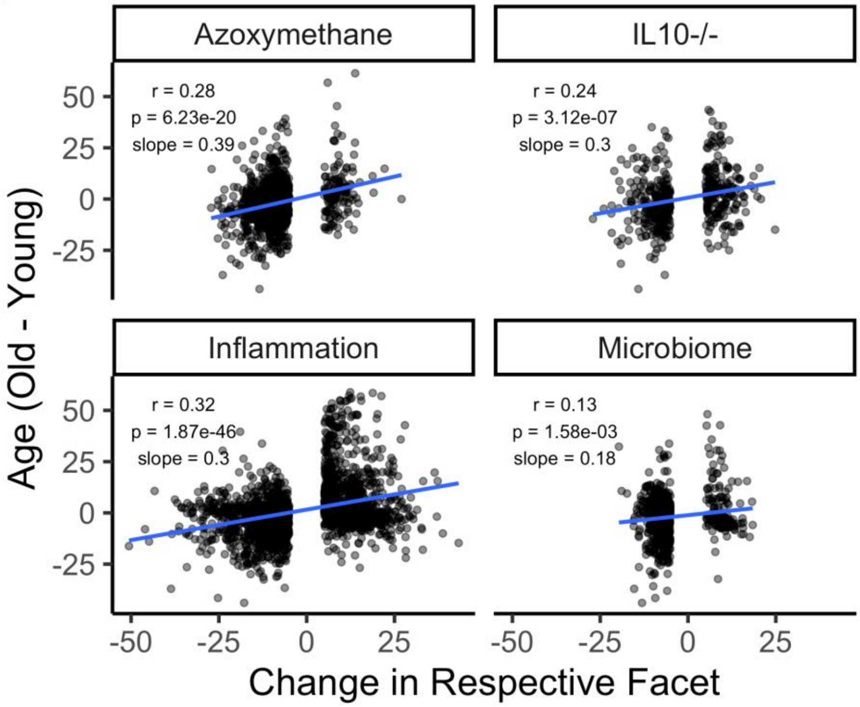
Azoxymethane
IL10-/-
Inflammation
Microbiome
r = 0.28
p = 6.23e-20
slope = 0.39
r = 0.24
p = 3.12e-07
slope = 0.3
r = 0.32
p = 1.87e-46
slope = 0.3
r = 0.13
p = 1.58e-03
slope = 0.18
50
25
0
-25
-50
Age (Old - Young)
Change in Respective Facet
